# A role for the immune system-released activating agent (ISRAA) in the ontogenetic development of brain astrocytes

**DOI:** 10.1101/2021.03.01.433338

**Authors:** Aminah M. I. Al-Awadi, Abdulaziz Isa AlJawder, Alyaa Mousa, Safa Taha, Moiz Bakhiet

## Abstract

The Immune System-Released Activating Agent (ISRAA) was discovered as a novel molecule that functions as a mediator between the nervous and immune systems in response to a nervous stimulus following an immune challenge. This research investigated the role of ISRAA) in promoting the ontogeny of the mouse brain astrocytes. Astrocyte cultures were prepared from two-month-old BALB/c mice. Recombinant ISRAA protein was used to stimulate astrocyte cultures. Immunohistochemistry and ELISA were utilized to measure ISRAA and IFN-γ levels, IFN-γR expression and STAT1 nuclear translocation. MTT-assay was used to evaluate cellular survival and proliferation. To assess astrocyte cell lysates and tyrosine-phosphorylated proteins, SDS-PAGE and western blot were used. ISRAA was highly expressed in mouse embryonic astrocytes, depending on cell age. Astrocytes aged seven days (E7) showed increased proliferation and diminished differentiation, while 21-day-old (E21) astrocytes depicted reversed effects. ISRAA stimulated the tyrosine phosphorylation of numerous cellular proteins and the nuclear translocation of STAT1. IFN-γ was involved in the ISRAA action as ISRAA induced IFN-*γ* in both age groups, but only E21 astrocytes expressed IFN-*γ*R. The results suggest that ISRAA is involved in mouse brain development through the cytokine network involving IFN-*γ*.

## Introduction

The novel immune modulator Immune System-Released Activating Agent (ISRAA) was recently discovered as a 125 amino acid protein of a 15kDa molecular mass. It is a protein secreted due to a central nervous system stimulus triggered by parasitic challenge and has been found to modulate the immune system by acting on the spleen (1). In the mouse system, ISRAA is located in intron 6 of the zmiz1 gene. Analysis of protein sequences has demonstrated that Fyn is ISRAA’s binding partner. *In vitro* studies using Fyn have shown that ISRAA is phosphorylated at tyrosine 102. In mouse splenocytes, ISRAA has been shown to downregulate T-cell activation and the phosphorylation of tyrosine Y416 in the Src-family kinases. Thus, ISRAA is considered to be a novel T-cell signalling molecule, induced by the nervous system (2). In biological assays, ISRAA illustrates dose-dependent biological activities, as low doses of 50pg/ml result in cell activation, while high doses of 5 μg cause apoptosis (3, 4). Analysis of green fluorescent protein (GFP) expression highlights that ISRAA is transcribed and expressed in lymphoid organs. Down regulation of T-cell activation and cytokine production has also been noted in the splenocytes of ISRAA knockout (-/-) mice, which show lymphoid organ hyperplasia. Furthermore, the transcription factor ELF1, which is involved in T cell regulation, is a binding partner for ISRAA, and a transcriptomic study using the ISRAA-overexpressing cell line EL4 has demonstrated the differential expression of numerous genes required for T-cell activation and growth (5).

Several studies have concluded that immune mediators, such as cytokines and chemokines, regulate the development and differentiation of embryonic brain cells. For example, the cytokine IFN-γ has been shown to be a factor in embryonic brain development (6) and in neural precursor regulation (7). It exhibits anti-proliferative effects, produces higher cell survival and induces major histocompatibility complex (MHC) expression in embryonic brain cells. Co-culture with the cytokine IL-4 abrogates the production of IFN-γ and blocks its anti-proliferative action (8). Also, the natural induction of a broad spectrum of cytokine profiles in first trimester human forebrain cells has been demonstrated at the gene level at five weeks’ gestation and elevates with age (9). Concerning chemokines, Rantes has been found to be an intrinsic immune mediator, playing a substantial role in brain ontogenetic development, and IFN-γ is crucially involved in this activity; the two mediators act as original signalling molecules in an effective network of cytokines and chemokines (10).

This research presents further evidence of the interaction between immune mediator networks – exemplified by ISRAA, IFN-γ and embryonic astrocytes – for possible ISRAA activity in astrocyte survival and proliferation during brain development. It also considers the distinct signalling pathway ISRAA uses to perform its action.

## Materials and Methods

### Animals

Two-month-old BALB/c mice (30–40g) were used. Seven-day-old (E7) and 21-day-old (E21) embryos were obtained from 24 (12 per experiment per each time point) pregnant mice. Ethical approval was obtained from the Ethics Committee of the Medical College of the Arabian Gulf University (AGU), Bahrain. Experiments were accomplished via relevant guidelines and regulations. Animal experiments were carried out under the Guide of the National Institute of Health for the Care and Use of Laboratory Animals (NIH Publications No. 80-23, revised 1996). The animals were housed four per cage, given water ad libitum, provided with food and maintained on a 12h/12h light/dark cycle until they were sacrificed. All efforts were made to decrease the number of animals used and the suffering of those sampled.

### Astrocyte culture

Pregnant Balb/c mice of gestational age 7 days (E7) and 21 days (E21) were killed by cervical dislocation, and uteri were aseptically removed and transferred to Petri dishes containing sterile PBS (Sigma). Each decidua from the uterine sac was dissected and transmitted to a new sterile Petri dish containing PBS to rinse away excess blood. Deciduas were then transferred to another sterile Petri dish containing PBS, and the amniotic sac was gently removed. Embryos were extracted from the amnion; the heads were then dissected. The brains removed into another Petri dish containing PBS, and crushed through a 70μm nylon mesh. Astrocytes were then prepared as previously described (11). In brief, glial cells were suspended in Dulbecco’s Modified Eagle’s Basal Medium (Fisher Scientific, Roskilde, Denmark) and placed in 50cm^2^ flasks at 10^6^ cells/ml. The medium was complemented with 2mM glutamine, 50 μg/ml streptomycin, 50IU/ml penicillin, 10mM Hepes and 1% MEM amino acids (all obtained from Merck, Darmstadt, Germany). Cells were grown for two weeks. Eight to ten days later, microglia were growing on the surface, and the astrocytes were detached by shaking for 2h at 260 rounds per minute (RPM). Monolayers of astrocytes in 50cm^2^ flasks were given a final shaking at 100RPM for one day to remove any existing oligodendrocytes. Astrocyte purity was approximately 98%, as confirmed by the anti-glial fibrillary acid protein (anti-GFAP) antibody (Merck, Darmstadt, Germany).

### ISRAA protein preparation

The recombinant ISRAA protein was obtained as previously described (1).

### Cell cultures, ISRAA treatment and blocking experiments

Cells from E7 and E21 embryos were kept on polystyrene vessel tissue culture glass slides with four chambers (BD Falcon Tubes; BD Biosciences, Bedford, MA, USA), at a concentration of 1 in 0.5ml of Dulbecco’s Modified Eagle’s Basal Medium per chamber, for a total of 15 slides. Two chambers of each slide were treated with ISRAA at a concentration of 50pg/ml. This concentration was selected after performing titration study. Cells were then grown for ten days in a 37° C incubator with 5% CO_2_. Media and growth were checked every two days until the time of staining. The recombinant ISRAA used in the experiments was endotoxin free, and it was neutralised with Polymyxin B (Merck, Darmstadt, Germany). Blocking experiments used the anti-ISRAA antibody (1), the rat anti-mouse anti-IFN-γ antibody (Bio-Rad, Oxford, UK) and tyrphostin A47 (a tyrosine protein kinase [TPK] specific blocker; Merck, Darmstadt, Germany). For each antibody, a concentration of 5 μg/ml was employed, and 10^−6^M was used for tyrphostin A47.

### Immunohistochemistry

This technique was achieved as previously described (12). In brief, non-stimulated cells were washed in phosphate-buffered saline (PBS) three times. Cells were then blocked by 2% normal sheep serum diluted in PBS for 30min, followed by the addition of the rabbit anti-mouse IFN-γ receptor (IFN-γR) antibody (MyBioSource, Inc., San Diego, CA, USA) and the rabbit anti-ISRAA polyclonal antibody (1), diluted at 1/1,000, for 24h at 4°C. Biotinylated secondary anti-rabbit IgG antibody (Vector Lab., Burlingame, CA, USA) was added over the cells for 1h. DAB was used for staining (Vector Lab., Burlingame, CA, USA). Counter staining was conducted with haematoxylin for 15s, and slides were mounted with glycerol oil.

### MTT assay

MTT assay was used to detect the number of viable cells. Astrocytes from E7 and E21 embryonic brains were counted and cultured in 96-well plates mounted with poly-D-lysine (2.5×10^4^ astrocytes in 200 μl culture medium). Treatment with recombinant ISRAA and blocking tests were completed as described above. FCS was washed with RPMI; then, 100 μl of 1mg/ml MTT (Merck, Darmstadt, Germany) in RPMI 1640, without phenol red, was added. Plates were incubated at 37°C for 3h in a humidified, 5% CO_2_ incubator. The MTT reaction was stopped by adding 200 μl of 2-propanol for 30min. This was followed by vigorous mixing to assure the complete solubilisation of the dye using a multichannel pipette. An Enzyme-Linked Immunosorbent Assay (ELISA) reader (mcc/340, Labsystem, Helsinki, Finland) was used to quantify the amount of converted MTT at 540nm.

### Cell-Released Capturing (CRC)-ELISA for IFN-γ detection

To detect IFN-γ, CRC-ELISA was used (13). Briefly, flat-bottomed EIA/RIA plates (Daigger Scientific, Vernon Hills, IL, USA) were coated with 100 μl of rat anti-mouse IFN-γ mAb (10μg/ml; MyBioSource, Inc., San Diego, CA, USA). These capturing antibodies were diluted in carbonate–bicarbonate buffer of pH 9.6 and kept overnight at 4°C. Then, cells were washed four times with 0.05M Tween-PBS and blocked with 5% bovine serum albumin (BSA) for 90min at room temperature (RT). 200 μl suspensions of astrocytes were subsequently added (final concentration: 10^5^ cells/well). Some cultures were stimulated with 50pg/ml recombinant ISRAA, while others were kept without stimulation. Following a 48h incubation period in a humidified atmosphere of 7% CO_2_ at 37°C, astrocytes were removed with trypsin and washed Three times in PBS. Captured IFN-γ was detected by adding biotinylated goat anti-rat IgG antibody (Vector Lab., Burlingame, CA, USA), diluted 1:2,000 in PBS with 2% BSA and 0.5% Tween 20. This was followed by incubation for 1h at 37°C and three washes. Then, 100μl of avidin-biotin alkaline phosphatase complex (ABC-AP; Vector Lab., Burlingame, CA, USA), diluted in PBS (1:100), was added for 30min. This was followed by three washes with PBS to remove any unbound ABC-AP and the addition of 100µl/well of the enzyme substrate solution. Plates were incubated in the dark for 15min, and, thereafter, the absorbance was measured in a Multiskan photometer (mcc/340; Labsystem, Helsinki, Finland) at 405nm. Concentrations of the secreted IFN-γ were measured from a standard curve obtained in parallel, using known concentrations of recombinant mouse IFN-γ (Bio-Rad, Oxford, UK) and a capturing of anti- IFN-γ mAb, as previously detailed (13).

### Astrocyte cell lysate preparation

Aliquots of astrocyte lysates, each containing about 10^6^ cells/ml, were centrifuged at 800 x g for 10min at RT in 1.5ml Eppendorf tubes. Supernatants were removed, and cell pellets were suspended in 100µl of lysis buffer containing 137mM NaCl, 20mM TRIS-HCl (pH 7.5), 0.5% sodium deoxycholate, 0.1% SDS, 1% Triton X-100, 1mM phenylmethylsulphonyl fluoride, 10% glycerol, 2mM EDTA, 0.15U/ml aprotinin, 25mM β-glycerophosphate and 1Mm sodium orthovanadate. Cell suspensions were then lysed by shock freezing in liquid nitrogen and kept at -70°C.

### SDS-PAGE and western blot assay

SDS-PAGE and western blot assay were used for astrocyte cell lysates and for tyrosine-phosphorylated proteins. 30 μl of astrocyte lysates with equal protein content in 2× Laemmli buffer (10% glycerol, 2% SDS, 5mM EDTA, 5% β-mercaptoethanol, 62.5mM TRIS-HCl [pH 8.8] and 10μg/mL bromophenol blue) were mixed and boiled for 3min and then centrifuged at 12,000 x g for 15s. Lysate concentrations were measured and adjusted to obtain equivalent amounts of protein before being loaded on a 10% polyacrylamide gel (30 μl of the mixtures having equivalent protein amounts). Ponceau S staining solution (Merck, Darmstadt, Germany) was used to confirm protein loading equivalence. Following SDS-PAGE, electroblotting was employed to transfer proteins to a nitrocellulose membrane. Unspecific binding sites were blocked with PBS-Tween 20 (0.1%)+ 1% BSA for 1h at RT. Membranes were then incubated with 0.5 μg/ml of mouse anti-phosphotyrosine monoclonal antibody 4G10 (Merck, Darmstadt, Germany) at 4°C overnight. After washing three times in PBS-Tween 20 (0.1%), blots were incubated with rabbit anti-mouse IgG peroxidase-conjugated, diluted at 1:5,000 (Merck, Darmstadt, Germany) for 90min at RT and developed with the enhanced chemiluminescence (ECL) system (Bio-Rad, Oxford, UK). To control the specificity of the identified phosphorylated protein, an irrelevant control antibody was used.

### STAT1 nuclear localisation

Immunofluorescence was used to detect the nuclear translocation of STAT1. A 1ml astrocyte cell suspension of 4×10^4^ cells/ml was plated in three chambers of four-chamber glass slides (BD Falcon, USA) coated with poly-D-lysine. 50pg/ml of recombinant ISRAA were added to the first well. Tyrphostin A47 (a protein kinase inhibitor), in the non-toxic concentration of 10^−6^M (14), was added (10 μl) to the second well. The third, non-stimulated well was used as a control. The slides were then incubated for 5min, 15min and 60min at RT. After rinsing twice in PBS for 20min, the cells were fixed for 2min in a methanol and acetone solution. Following two more washes in TBST (containing 10mM tris-Cl [pH 8.0] and 100mM NaCl, 0.02% Tween 20) the astrocytes were incubated in TBST containing 3% BSA for 40min. Anti-STAT1 primary antibody (rabbit polyclonal) (Abcam, Cambridge, MA, USA) was added at a final dilution of 1:100. Slides were then incubated overnight at 4° C. After being washed three times in TBST, the secondary antibody Cy^3^ (fluorescein-conjugated goat anti-rabbit IgG antibody; Abcam, Cambridge, MA, USA) was added in a final dilution of 1:200, followed by incubation for 70min at RT. Slides were then washed three more times in TBST for about 10min, and, thereafter, they were mounted with Kaiser glycerine gelatine (Merck, Darmstadt, Germany).

### Statistical analysis

Student’s two-sided, unpaired t-test was chosen to calculate the statistical significance between two groups. *P*<0.05 was considered significant in each test.

## Results

The results of this work indicate the role ISRAA plays in mouse brain astrocytes development, through the cytokine network involving IFN-γ as illustrated in the graphical abstract (Fig 1).

**Figure 1.**
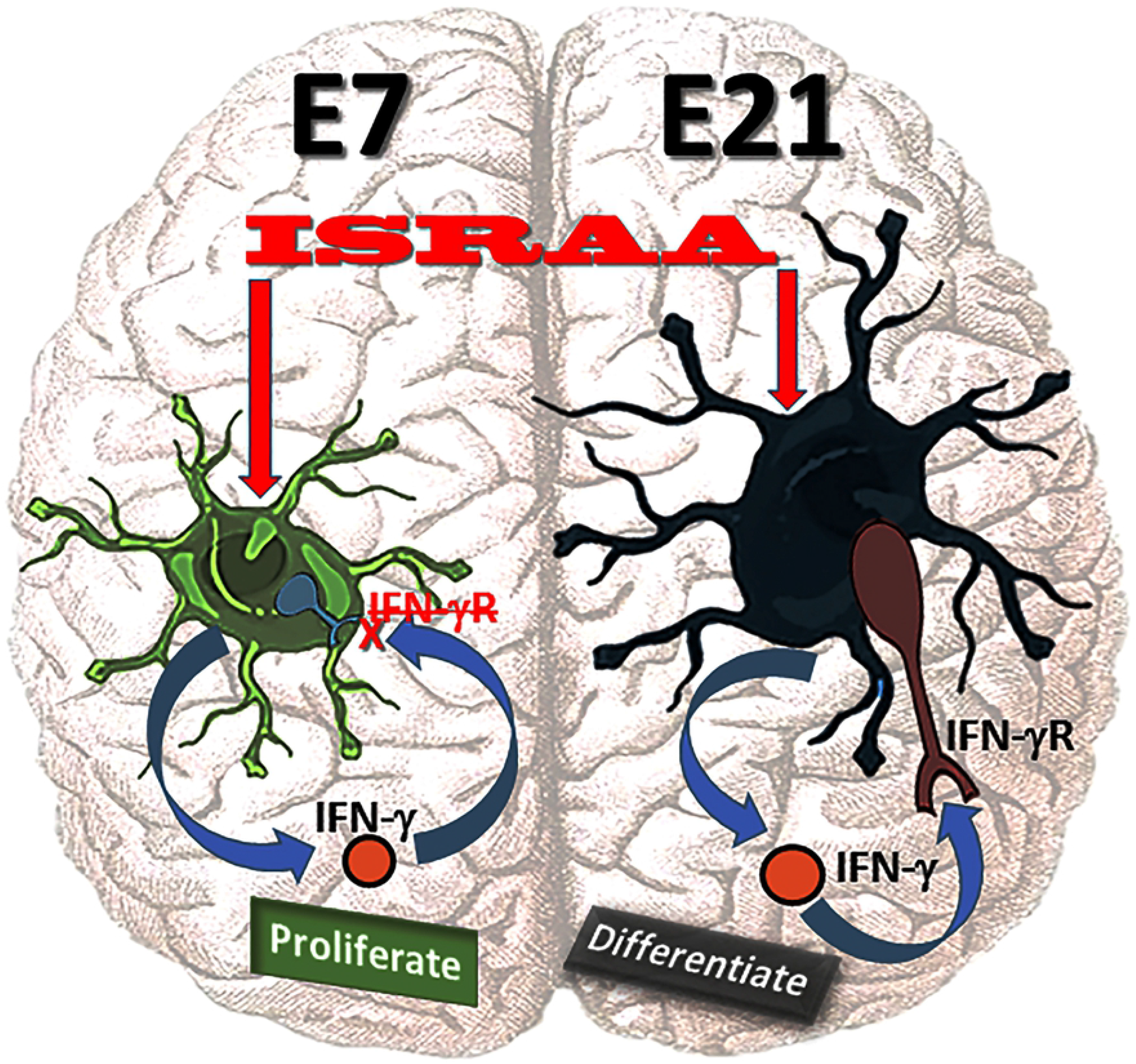
Graphical abstract illustrating the role of ISRAA in mouse brain astrocytes growth regulation through the cytokine network involving IFN-γ. ISRAA acted on embryonic brain astrocytes early at E7 and late at E21. It stimulated them to produce IFN-γ. E7 astrocytes did not express IFN-γR; therefore, they could not respond to the anti-proliferative effects of IFN-γ and continued to proliferate. However, E21 cells did express IFN-γR, and, accordingly, they ceased proliferation and began to differentiate.

To assess intrinsic ISRAA expression in embryonic mouse brain astrocytes, the researchers performed immunohistochemical staining. ISRAA immunopositive astrocytes were detected in elevated numbers in an age-dependent manner, as E7 astrocytes showed few faint cells stained for ISRAA (Fig 2A), while E21 cells expressed high ISRAA levels with strong staining (Fig 2B).

**Figure 2.**
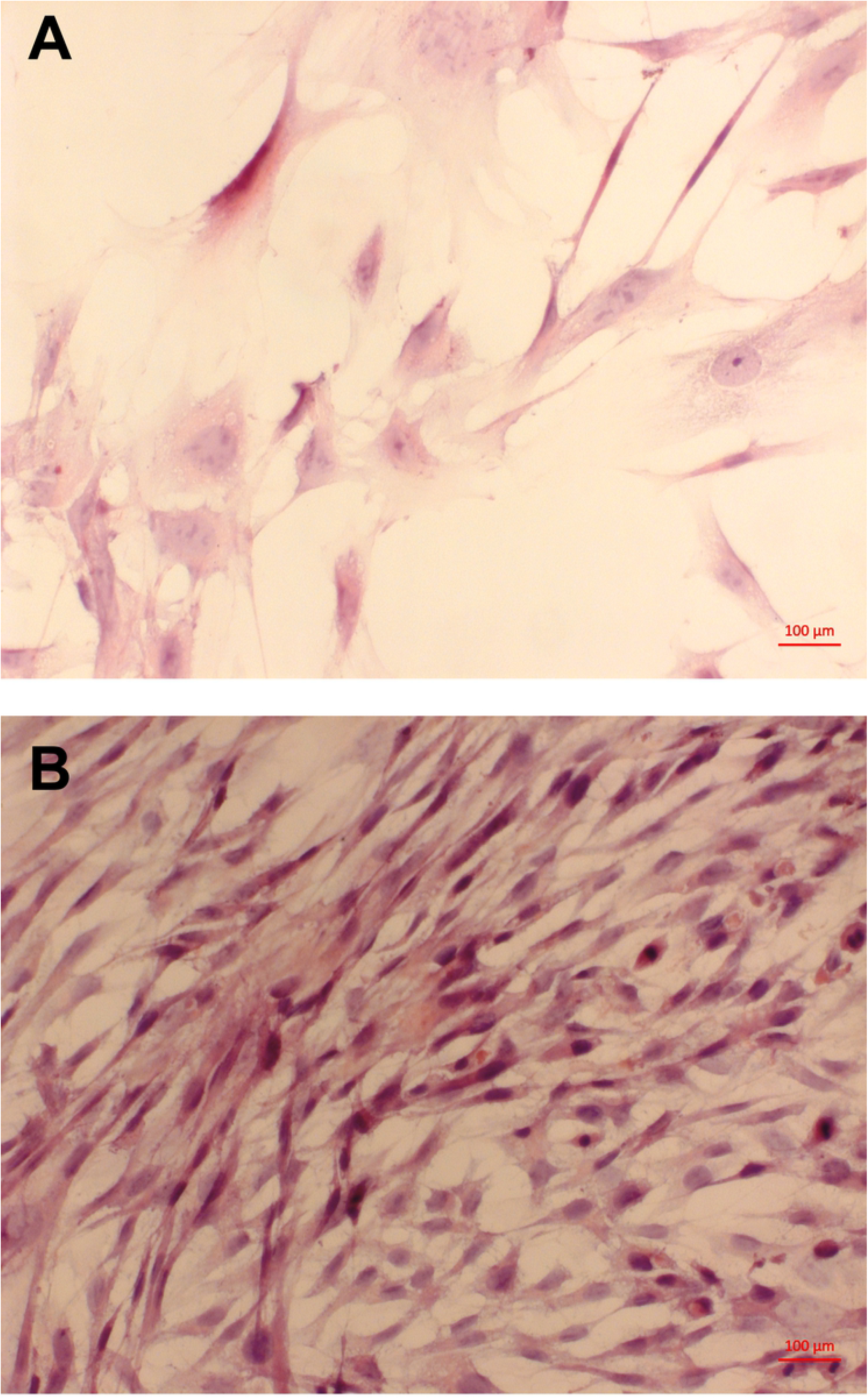
Illustration of ISRAA expression by embryonic brain astrocytes. Intrinsic ISRAA expression in embryonic mouse brain astrocytes was detected by immunohistochemistry. Ten fields were counted in each chamber of the tissue culture glass slides containing astrocytes from E7 and E21. (A) demonstrates few faint cells stained for ISRAA in E7. However, E21 showed significantly (*p*<0.05) high numbers of positively and strongly stained cells for ISRAA (B). Pictures were acquired via light microscopy (x630).

The capacities of the embryonic mouse brain astrocytes to proliferate and to differentiate were assessed via MTT assay and morphological determination, respectively. Non-stimulated E7 astrocytes exhibited higher proliferative responses without obvious differentiation (Fig 3A and Fig 4A), while E21 astrocytes showed clear cell differentiation, as well as lower proliferation, compared to E7 astrocytes (*p*<0.05) (Fig 3A and Fig 4B).

**Figure 3.**
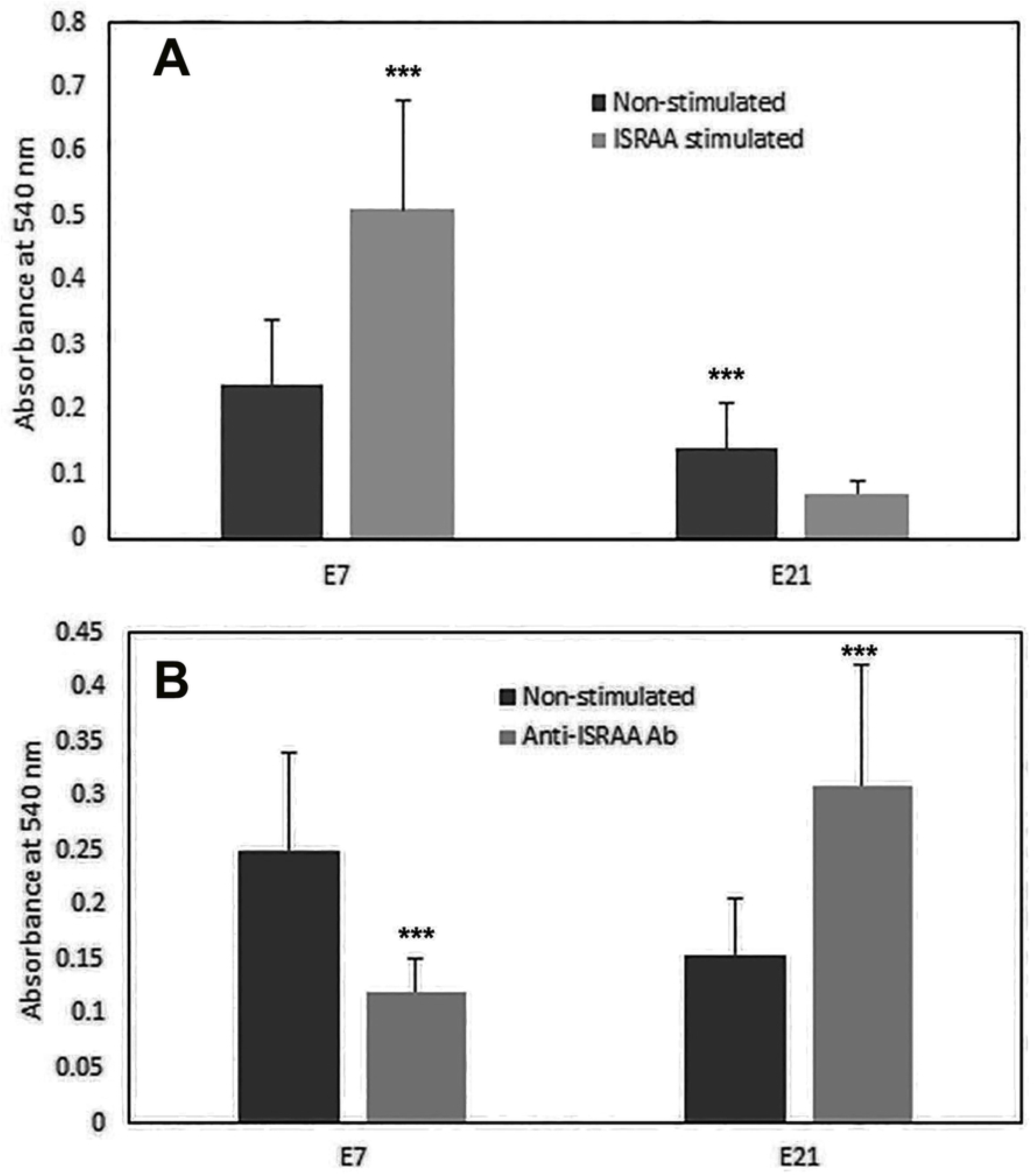
Assessment of ISRAA induced proliferation and anti-ISRAA antibody blocking effects. Cell proliferation was measured via MTT. Non-stimulated E7 astrocytes showed higher proliferative responses, while E21 astrocytes exhibited lower proliferation compared to E7 astrocytes (\*\*\**p*<0.05). E7 showed increased cell proliferation in response to recombinant ISRAA (\*\*\**p*<0.05). However, less proliferation was detected in E21 when these cells were stimulated with ISRAA (Fig 3A). Pre-incubation with anti-ISRAA antibody abrogated the proliferative response in E7 and induced E21 to proliferate (\*\*\**p*<0.05) (Fig 3B). Experiments were conducted twice with alike findings. Bars represent the means ± SDs of eight cultures.

**Figure 4.**
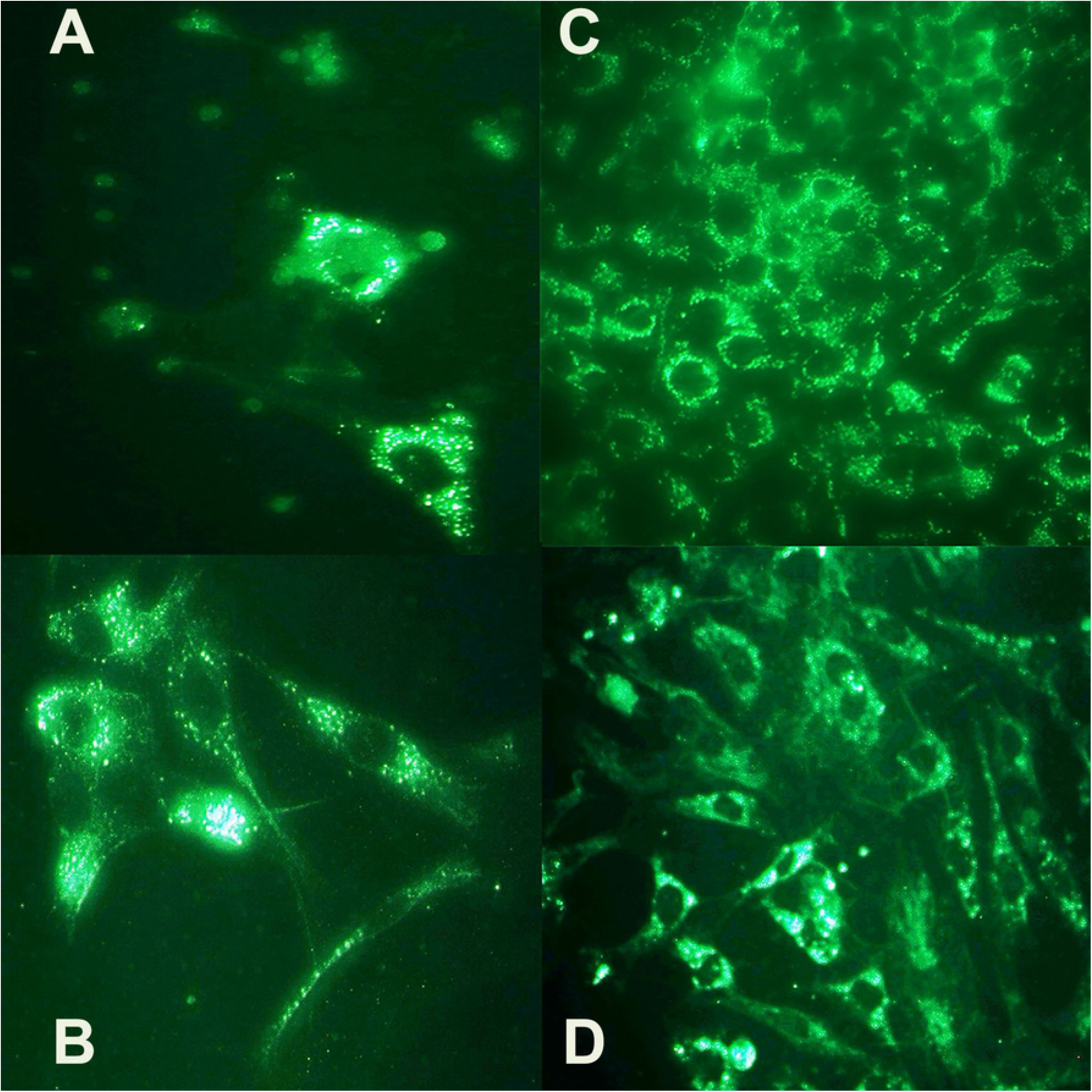
Embryonic astrocyte proliferation and differentiation. Spontaneous and ISRAA induced cell proliferation and differentiation were demonstrated by morphological observation and determination. Astrocytes were prepared and cultured as described in the Materials and Methods section. (A) demonstrates non-stimulated astrocytes from E7 with high proliferation and no clear differentiation. E21 astrocytes showed clear cell differentiation with lower proliferation compared to E7 (B). Stimulation of E7 with recombinant ISRAA resulted in increased cellular proliferation (C), while E21 showed well differentiated cells and less proliferation in response to ISRAA stimulation (D). Pictures were gathered via bright field microscopy (x630). (Statistical significance is shown in Fig 3).

The responses of E7 astrocyte cultures to the addition of recombinant ISRAA resulted in enhanced cellular proliferation (Fig 3A and Fig 4C) (*p*<0.05), while E21 astrocytes showed less proliferation, with more well differentiated cells (Fig 3A and Fig 4D). Interestingly, incubation with the anti-ISRAA antibody suppressed the proliferative response in E7 astrocytes but induced proliferation in E21 astrocytes (*p*<0.05) (Fig 3B).

To examine the potential process by which ISRAA persuades diverse activities at different developmental stages, the researchers studied IFN-γ induction after ISRAA stimulation. The rationale behind this experiment was that IFN-γ has been previously shown to have an anti-proliferative effect on embryonic cells (9, 10). Both E7 and E21 astrocytes showed spontaneous IFN-γ production, which was significantly increased after ISRAA stimulation (*p*<0.05). IFN-γ levels were found to be significantly elevated in E21, compared to E7, astrocyte cultures (*p*<0.05) (Fig 5).

**Figure 5.**
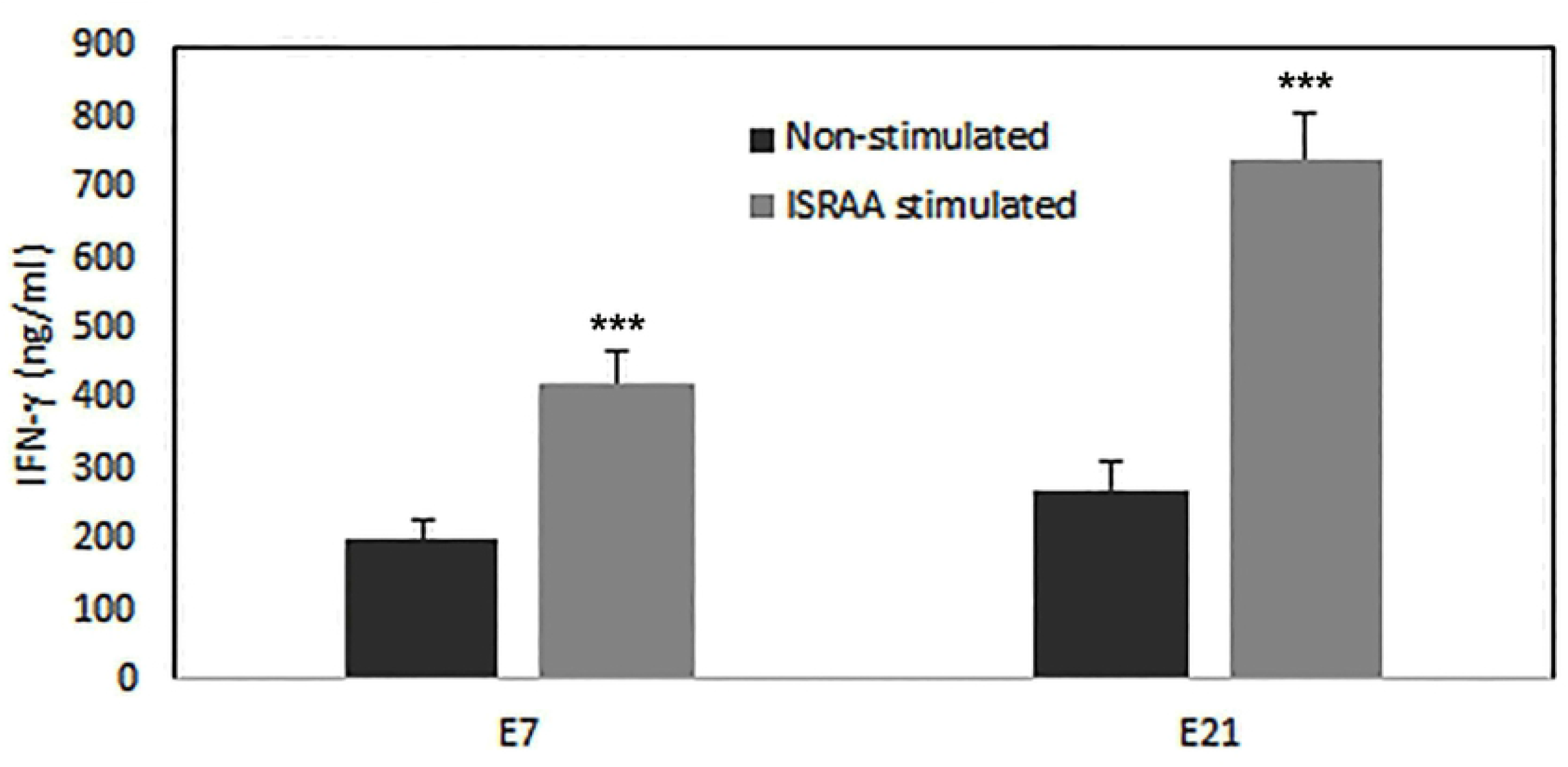
Spontaneous and ISRAA-induced IFN-γ production by embryonic astrocytes. E7 and E21 astrocytes spontaneously produced IFN-γ. Stimulation with recombinant ISRAA significantly increased IFN-γ production as measured by CRC-ELISA (see Materials and Methods) (\*\*\**p*<0.05). The induction of IFN-γ was significantly higher in E21 than in E7 astrocyte cultures (\*\*\**p*<0.05). Experiments were conducted twice with alike findings. Bars represent the means ± SDs of 12 cultures.

Excitingly, IFN-γR was not expressed in E7 astrocytes (Fig 6A) but was expressed in E21 astrocytes (Fig 6B). Blocking of IFN-γ with the anti-IFN-γ antibody in E21 astrocyte cultures inverted the inhibitory action of ISRAA on proliferation, but it had no effect on E7 cultures (Fig 7).

**Figure 6.**
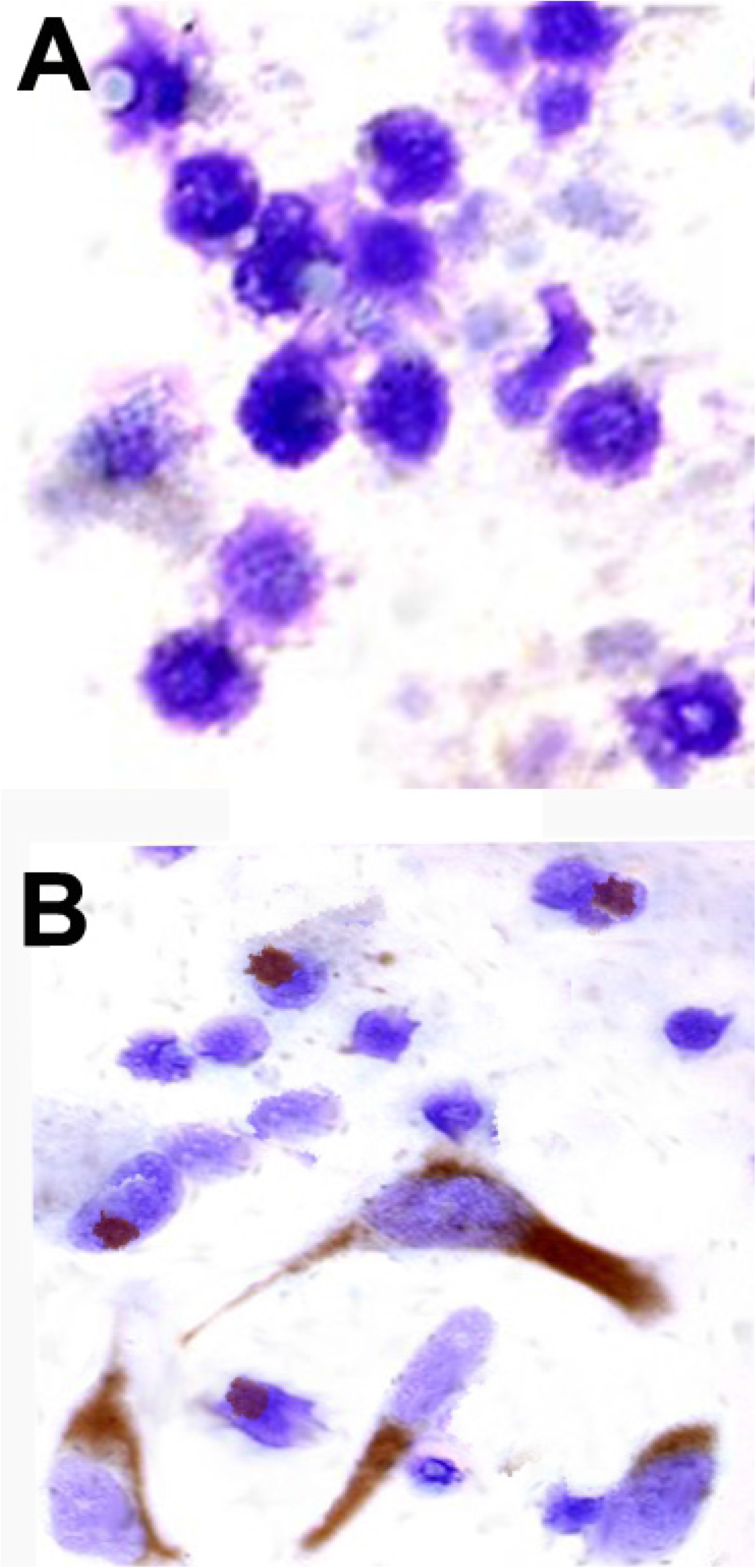
Expression of IFN-γR by embryonic astrocytes. As demonstrated here, E7 astrocytes did not express receptors for IFN-γ (A), while E21 astrocytes showed strong expression of IFN-γR (Fig 6B). Pictures were acquired with light microscopy (x630). Ten fields were counted in each chamber of the tissue culture glass slides containing astrocytes from E7 and E21. The number of cells expressing IFN-γR at E21 was significantly higher than that at E7 (p<0.05).

**Figure 7.**
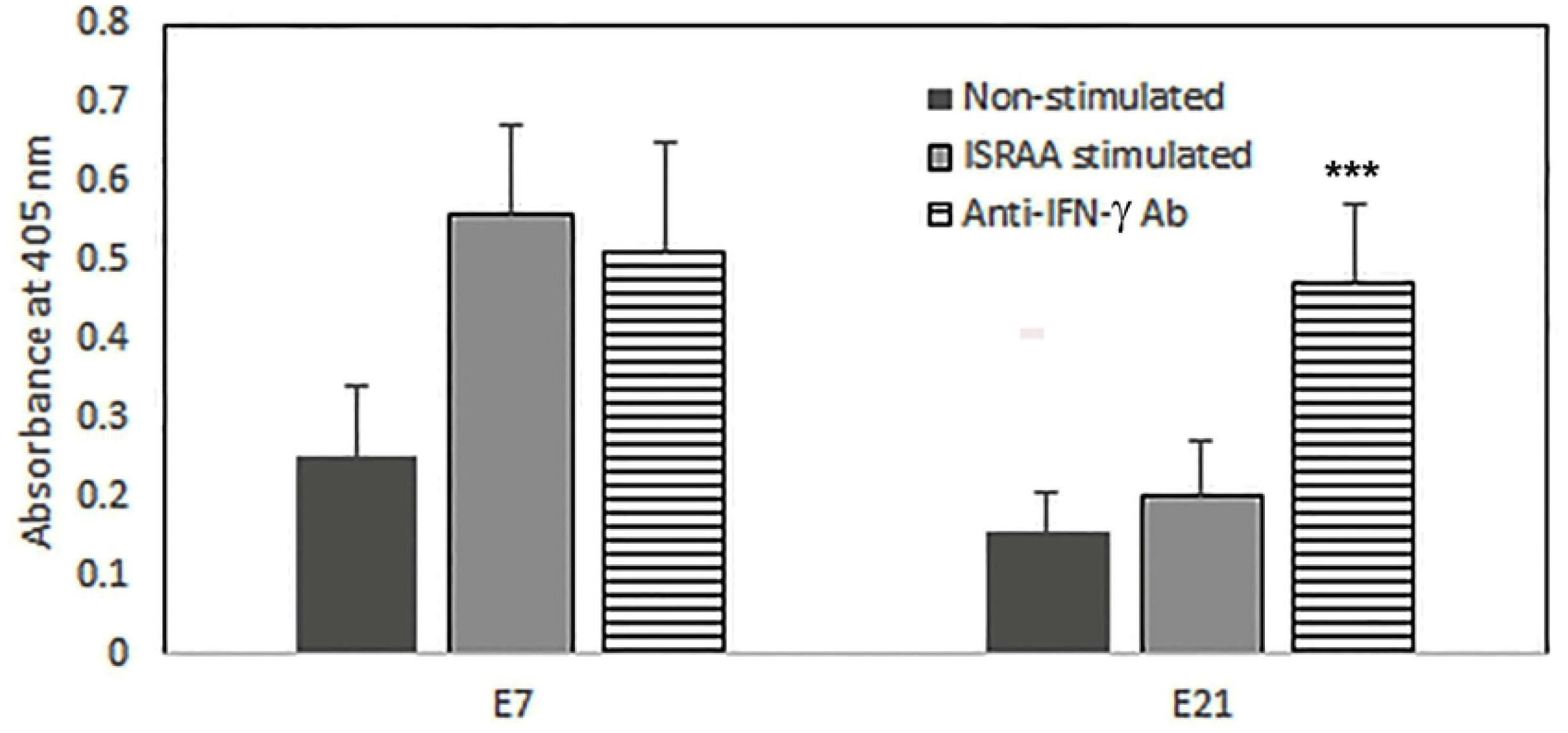
Blocking results of anti-IFN-γ antibody on ISRAA proliferation suppression at E21. Incubation with anti-IFN-γ antibody in cultures of E21 astrocytes significantly inverted the blocking effects of ISRAA on proliferation (***p<0.05) but had no effect on E7 (bars with horizontal lines). Experiments were conducted twice with alike results. Bars represent the means ± SDs of 12 cultures.

Western blot analyses were used to study tyrosine kinase activity in the embryonic astrocytes after ISRAA stimulation by analysing cell lysates from E7 astrocytes prepared 5min, 15min and 60min after ISRAA stimulation. Small numbers of tyrosine phosphorylated proteins were demonstrated in the non-stimulated, control lysates. However, a strong and rapid elevation in tyrosine phosphorylation was noted after ISRAA stimulation; it reached a high level after 15min and decreased to control levels after 60min. Incubation with the tyrosine kinase specific inhibitor tyrphostin reduced ISRAA activity on protein kinase expression at the various time points (Fig 8). The induction of tyrosine phosphorylation noted in E21 astrocytes was similarly observed in E7 astrocytes.

**Figure 8.**
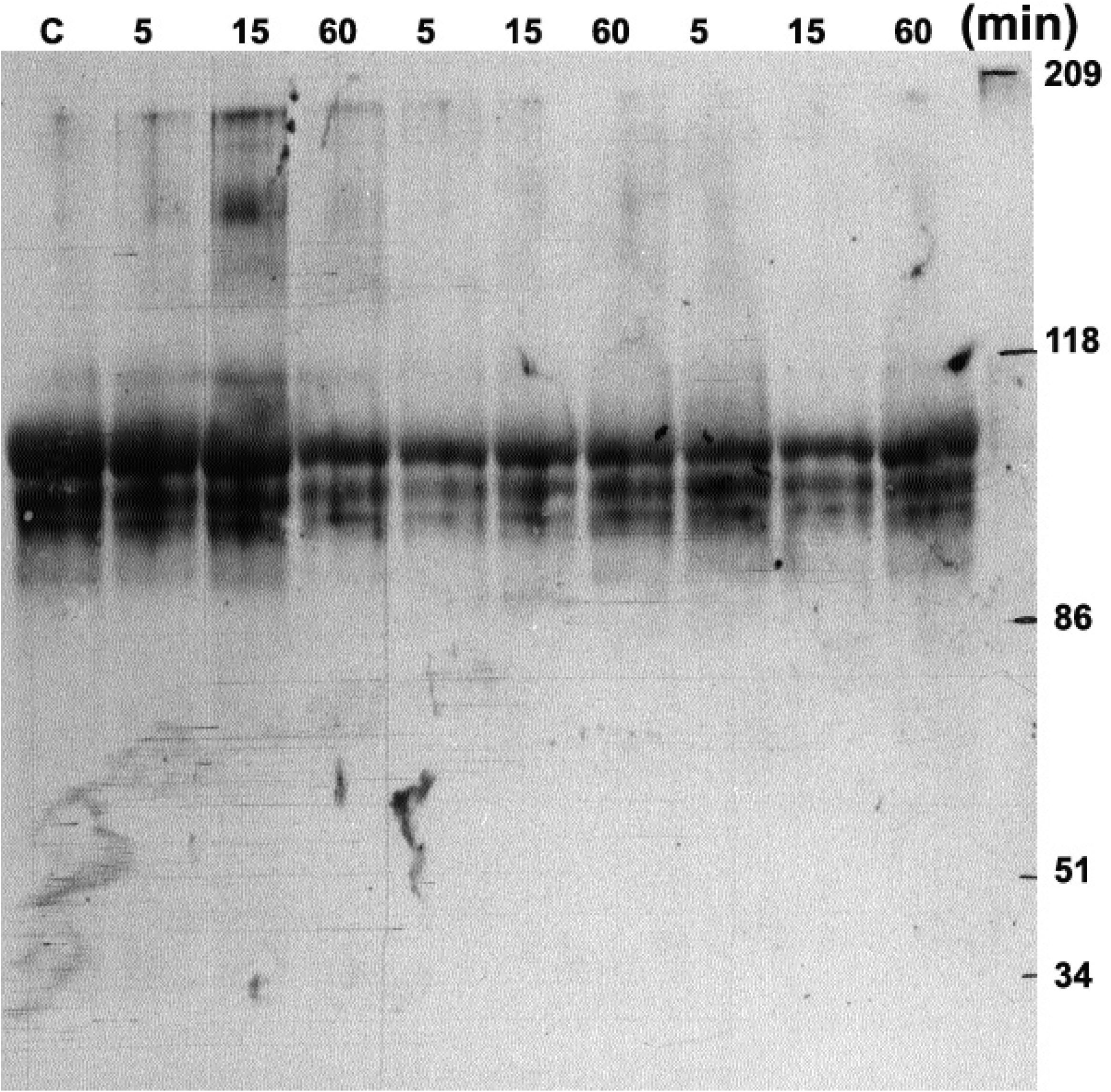
Phosphorylation of tyrosine-specific protein. Tyrosine kinase activity in E7 astrocytes after ISRAA stimulation was examined using western blot analyses. Cell lysates were prepared 5min, 15min and 60min after ISRAA stimulation. Note the slight amounts of tyrosine phosphorylated proteins in the non-stimulated control lysates (C) on lane 1. ISRAA stimulation resulted in a strong and fast rise in tyrosine phosphorylation, which reached its highest point by 15min and declined to the control level at 60min. An irrelevant antibody was used to control the specificity of the phosphorylated protein. The band representing activated TPKs in lane three (middle lane) was not detected. Tyrphostin A47 (a tyrosine kinase specific inhibitor) blocked the ISRAA induced protein kinase at the various detection times. Inhibition of the strongest activity at 15min is demonstrated here.

The localisation of the STAT1 transcription factor in E7 astrocytes stimulated with ISRAA was examined via a fluorescent antibody. The anti-STAT1 antibody showed a generalised cytoplasmic antigen distribution prior to ISRAA treatment (Fig 9A) and a prominent nuclear fluorescence after 15min of ISRAA stimulation (Fig 9B). The translocation of STAT1 was prevented by the TPK specific inhibitor tyrphostin A47 upon co-culturing with the astrocytes (Fig 9C). The kinetics of STAT1 translocation induced by ISRAA were examined by tracking nuclear translocation for 5min, 15min and 60min. At the 5min mark, approximately 20% of the cells already depicted STAT1 nuclear translocation. This reached 95% at 15min, and then decreased to less than 5% at 60min. The nuclear translocation was demonstrated by positive nuclear immunostaining. Less than 1% of non-stimulated cells and cells treated with TPK specific inhibitor tyrphostin A47 showed positive nuclear translocation at 15min. Similar translocation of STAT1, found in E7 astrocytes, was also registered in E21 astrocytes.

**Figure 9.**
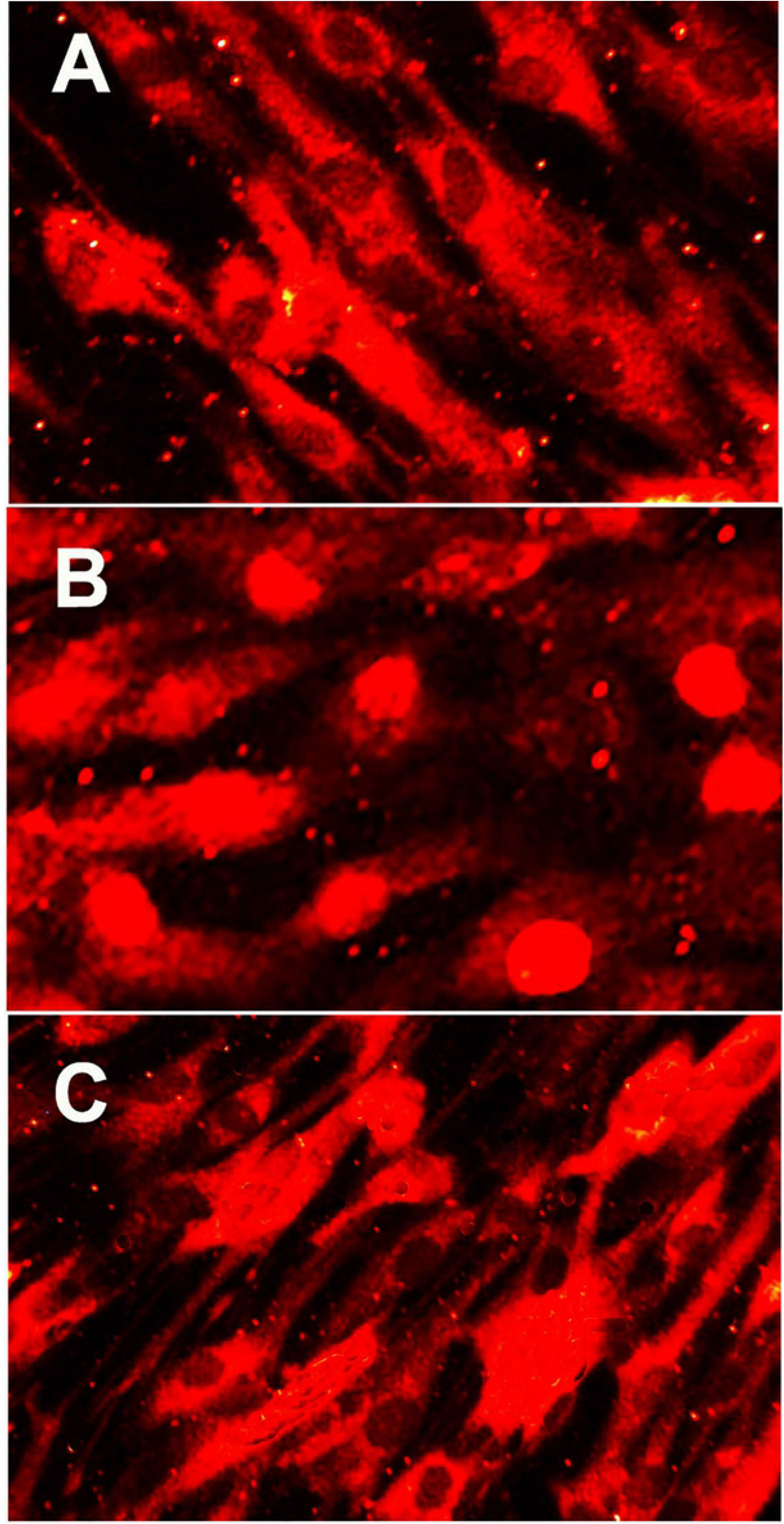
Nuclear localisation of the STAT1. Nuclear localisation of the STAT1 protein in E7 astrocytes stimulated with ISRAA. Pictures were obtained by a digital camera using a fluorescence microscope at x630 magnification after 15min incubation. (A) represents non-stimulated cells (note the generalised cytoplasmic antigen distribution before ISRAA treatment), while (B) depicts the nuclear translocation of STAT1 upon ISRAA stimulation. (C) demonstrates the blocking effects of tyrphostin A47 on the ISRAA promoted nuclear translocation of STAT1. Similar translocation of STAT1 was also noted in E21 astrocytes.

## Discussion

The immune mediator ISRAA has been well identified as an immune modulator, which mediates signals between the nervous and immune systems (1, 2, 5). The IFN-γ cytokine is a major effector molecule in the immune system, and it has been shown to play a key role in regulating brain growth (10, 15). The present work has demonstrated the potential role of ISRAA and IFN-γ in the development of embryonic mouse brain astrocytes. The data shows that these mediators act in an age-dependent manner. For example, ISRAA induced E7 and E21 astrocytes to proliferate and secrete IFN-γ. However, IFN-γ showed no effects in E7 astrocytes as these cells did not express IFN-γR. As they had progressed in age during the early embryonic stages, E21 astrocytes expressed IFN-γR and could respond to IFN-γ, which resulted in ceased proliferation, allowing time for differentiation. The network of these mediators definitely involves several cytokines and chemokines, as previously shown for Rantes (CCL5) (10). Furthermore, a well-balanced level of all types of I IFNs has been found to be essential for healthy brain physiology, and dysregulation of this cytokine system may lead to brain interferonopathies (16).

Astrocytes are well known for playing critical roles in maintaining neuronal function in the central nervous system (CNS) by producing cytokines, chemokines and growth factors. Additionally, the gene expressions of numerous cytokines and chemokines are differentially regulated by the NF-kB signalling pathway during physiological and inflammatory situations in human astrocytes (17). Similarly, the authors have demonstrated the importance of cytokine receptors and the intracellular signalling pathway, as illustrated by IFN-γR and STAT1 activation in this study.

The role of IFN-γR is based on its maturation during development. Previous work among C57Bl/6 mice, through several developmental stages, has revealed that cytokines and cytokine receptors are expressed in different areas of the brain at different developmental stages. For instance, TGFβ2, TGFβR1 and TGFα are expressed in different brain regions at E15. Also, CSF1R is expressed by macrophages and microglia during the early stages of embryonic development at E11 and at E15, respectively (18). For IFN-γR, previous studies have shown that IFN-γR1 and IFN-γR2 are expressed in oocytes obtained from mice and in germinal vesicles and metaphase II stages from preimplantation embryos (19). However, this does not indicate the organ specific expression of IFN-γR and its specific role in organ development. This is considered in context with other cytokines and growth factors, which are expressed during this embryonic stage (20-22) but have not been detected in the brain until after E11 (18). The present study has found that E7 astrocytes lack IFN-γR and, consequently, that they can continue proliferating because they receive no blocking signals from the IFN-γ cytokine induced by ISRAA. E21 astrocytes did express IFN-γR, and, therefore, the proliferation blocking initiated by ISRAA was most probably conducted via the IFN-γ receptor; thus, differentiation was permitted. Accordingly, the age-dependent effects of ISRAA on astrocyte proliferation and differentiation suggests that ISRAA is essential to brain development, a supposition justified by the inhibitory effects of the anti-ISRAA antibody on astrocyte proliferation and differentiation. ISRAA was constitutively expressed, allowing it to stimulate or inhibit growth according to cell age. Most probably ISRAA acted in this way to regulate growth processes, as it enhanced astrocyte proliferation early, at E7, to promote cell expansion and, at the same time, stop their differentiation. Conversely, in E21 astrocytes, after growth and expansion, ISRAA stopped proliferation to promote cell differentiation.

It is certain that this activity was not solely dependent on one mediator; it involved a network of factors. With this in mind, the authors looked for the IFN-γ cytokine, which was previously demonstrated as suppressing proliferation and supporting the differentiation of embryonic astrocytes (8). IFN-γ was further found to promote the differentiation of embryonic cholinergic septal nuclei and the nearby basal forebrain (23). IFN-γ was constitutively produced and significantly enhanced in response to ISRAA stimulation, and its activity was determined to be dependent on its receptor’s maturation and expression, as E7 astrocytes did not express IFN-γR while E21 astrocytes did.

The authors further explored a signalling pathway for ISRAA in the early stages of brain growth. The resulting data showed that tyrosine kinases were activated in the ISRAA signalling system by tyrosine phosphorylation of JAK kinases, along with STAT1 protein tyrosine phosphorylation. This was followed by nuclear translocation to initiate the gene transcription required for either proliferation or differentiation. In agreement with this finding, IFN-γ was illustrated to regulate proliferation and neuronal differentiation via STAT1 (15), and IFN-γ inhibited the differentiation of neural progenitor cells (NPCs) by negatively regulating Neurog2 expression through the JAK/STAT1 pathway (6).

## Conclusions

In conclusion, the preceding data provide evidence that ISRAA and IFN-γ are potentially intrinsic developmental brain astrocytes factors. Therefore, understanding the role of such inflammatory mediators as brain development regulatory agents, as well as their action networks, is essential to planning future therapies for neuroinflammatory disorders. However, one limitation on this study is the lack of *in vivo* data to support the presented *in vitro* results. But, on the other hand, a detailed *in vivo* study using ISRAA knockout mice that were recently generated in our laboratory (5) is currently initiated to demonstrate the *in vitro* effects *in vivo* on brain slices to further define the effects of ISRAA on astrocytes.

## List of abbreviations

-/-: Knockout mice
ABC-AP: Avidin-Biotin Alkaline Phosphatase Complex
AGU: Arabian Gulf University
BSA: Bovine Serum Albumin
CNS: Central Nervous System
E21: Astrocytes aged twenty one days
E7: Astrocytes aged seven days
ELISA: Enzyme-Linked Immunosorbent Assay
FCS: Foetal Calf Serum
GFAP: Glial Fibrillary Acid Protein
GFP: Green Fluorescent Protein
ISRAA: Immune System-Released Activating Agent
MHC: Major Histocompatibility Complex
NIH: National Institute of Health
NPCs: Neural Progenitor Cells
PBS: Phosphate-Buffered Saline
RPM: Rounds Per Minute
RT: Room Temperature

## Declarations

### Ethics approval and consent to participate

Ethical approval was obtained from the Ethics Committee of the Medical College of the Arabian Gulf University (AGU), Bahrain. Consent to participate “Not applicable”.

### Consent for Publication

“Not applicable”.

### Data availability

Data and material are available upon request.

## Authors’ contributions

AA did experimental work, AI did experimental work, AM contributed to the confocal microscope experiments and the analysis, ST participated in the lab work and the data analysis, MB is the principle investigator of the project and wrote the paper.

## Acknowledgements

“Not applicable”.

